# Reusability and composability in process description maps: RAS-RAF-MEK-ERK signalling

**DOI:** 10.1101/2020.12.08.416719

**Authors:** Alexander Mazein, Adrien Rougny, Jonathan R. Karr, Julio Saez Rodriguez, Marek Ostaszewski, Reinhard Schneider

## Abstract

Detailed maps of the molecular basis of the disease are powerful tools for interpreting data and building predictive models. Modularity and composability are considered necessary network features for large-scale collaborative efforts to build comprehensive molecular descriptions of disease mechanisms. An effective way to create and manage large systems is to compose multiple subsystems. Composable network components could effectively harness the contributions of many individuals and enable teams to seamlessly assemble many individual components into comprehensive maps. We examine manually-built versions of the RAS-RAF-MEK-ERK cascade from the Atlas of Cancer Signalling Network, PANTHER and Reactome databases and review them in terms of their reusability and composability for assembling new disease models. We identify design principles for managing complex systems that could make it easier for investigators to share and reuse network components. We demonstrate the main challenges including incompatible levels of detail and ambiguous representation of complexes and highlight the need to address these challenges.

## 1. Introduction

Detailed descriptions of disease mechanisms on the level of molecular processes have recently become available [1,2], with many examples of practical applications in the field of cancer research [3–7]. These disease maps are needed for integrating scattered knowledge and for advanced data interpretation and hypothesis generation [1,2]. The information is stored in a standard format that is both human- and machine-readable. The development of such maps is currently a time-consuming process that often requires extensive collaboration among multiple researchers and reuse of existing map components.

Bringing together map components from different sources presents challenges related to the heterogeneity of map components. Currently, the composition of map components is done ad hoc. While it is possible to check compatibility and then modify incompatible parts for integrating them into the developing resource, such ad-hoc assembly is difficult to scale up. With each reuse, map components are likely to be modified. In particular, the current ad-hoc approach to composition makes it difficult for large teams of collaborators to work together to compose map components created by different team members into large, integrated maps.

In practical terms, even for well-described pathways that are accessible in high-quality pathway databases, while developing a new map, one would need to decide which of the available components (pathway or subpathway) they should reuse for their project. The same component is often repeated with certain modification in the same database, and criteria for selecting one version over another is not always clear. The briding elements that connect map component to other pieces (for example, shared proteins in particular posttranslational-modification states) could be represented in an inconsistent way and therefore composability might be questionable.

### In this work, we aim to identify specific challenges in reusing components of existing maps and to define guidelines for building composable network modules

We argue that ensuring composability could ease the collaborative assembly of disease maps. Composability here is a practical approach to use a minimal set of universally applicable components, an approach in which the step of modification of components to make them compatible with the other components is avoided. Indeed, applying this design principle would allow assembling maps from high-quality, self-contained, reusable components that are individually easy to build and update.

Community-driven disease model development requires sharing components and minimising overlapping work [1,2]. Modularity and composability as design principles have been previously discussed in connection to modelling in the Physiome project [8]. The ongoing COVID-19 Disease Map effort [9] demonstrates new challenges for fast-track development of large-scale reconstructions of disease mechanisms, as well as the importance of the required technologies for integrating components provided by different groups, verifying their quality and ensuring their compatibility.

To develop principles for modularly composing disease maps, we focus on the RAS-RAF-MEK-ERK signalling cascade, one of the most well-studied signalling pathways. This pathway regulates key cellular functions such as growth and differentiation, and it is one of the most commonly mutated pathways in cancer [10–13]. ERK protein can activate hundreds of proteins, is regulated by multiple mechanisms at different steps of the signalling cascade, and it includes negative feedback loops and temporal and spatial regulation via compartmentalisation through binding to scaffold or adaptor proteins such as KSR1 and SEF [14–16]. This allows selective phosphorylation of specific ERK substrates in a context-dependent manner [14]. Because of its relevance to multiple diseases and because of its complexity, it is challenging to describe the complete molecular details of this cascade. As a result, the RAS-RAF-MEK-ERK cascade is a favourable example for investigating the reusability and composability of components in network biology in general and in disease maps in particular.

Assembling modular disease maps from heterogeneous components requires components that are reusable, compatible and composable. To discuss related issues, we would like to briefly outline the terms of *modularity*, *reusability*, *compatibility* and *composability* in the context of process description maps [17,18]. To a certain extent, the meaning of these terms overlap but they cover different aspects of the issue. *Modularity* is a desired property of networks, and *reusability*, *compatibility* and *composability* are desired properties of their components. *Modularity* can be defined as the degree to which a network part (module) can be separated from its parent network. One benefit of modular networks is that their components can often be flexibly reassembled into new networks. *Reusability* is the ability to use existing components for building new maps. Potentially, reusable modules can be employed in a context other than the one they were initially developed for. The use of standards is an important enabler of reusability [17,18] and on the notation in CellDesigner consistent with an earlier version of the SBGN standard (http://celldesigner.org). *Compatibility* means that network components can coexist without producing any undesired effects. In process description networks, events can be represented in different ways and on different levels of granularity. *Composability* is the ability to assemble components/modules into larger networks in various combinations without the necessity to modify components to make them compatible. One benefit of modular composable components is that large networks can be improved over time by swapping components for more accurate versions.

For the purpose of this paper, we leave out of our discussion such related issues as quality and styles of curation. This paper focuses on a set of highly and consistently curated maps for which annotation is done according to the best curation practices with the use of compatible identifiers. For example, UniProt IDs are available for proteins, ChEBI IDs for metabolites, each protein modification is described properly and each complex composition is reflected in its content. The quality and consistency of their curation guarantee a minimal level of compatibility.

## 2. Results

To investigate the compatibility and reusability of descriptions of biochemical networks, we reviewed versions of the RAF-MEK-ERK pathway in three databases of high-quality (Supplementary Table S1), manually-curated representations of process description networks: the Atlas of Cancer Signalling Networks (ACSN) [19], PANTHER [20] and Reactome [21,22].

The Atlas of Cancer Signalling Networks represents the RAS-RAF-MEK-ERK events in several ways that are partially compatible with each other. By design, each component is compatible with the surrounding network in each of the maps in the ACSN but it is not necessarily fully cross-compatible with other maps within the ACSN, which means that one representation can not be replaced by another representation. This allows investigating composability issues while focusing only on specific components and not on the whole network. As soon as the components are interchangeable they would be considered composable and compatible with any of the maps within the ACSN. Three maps - Adaptive Immunity, Innate Immunity and Cancer-Associated Fibroblasts - contain the same canonical representation of the cascade (Figure 1a); the Cell Survival map includes both the canonical (generally accepted and repeated in different databases) representation of MEK and ERK activation, as well as their spatial regulation by SEF/IL17RD [14,23] (Figure 1c) and KSR1 scaffold mechanism via BRAF (Supplementary Figure S1); and the EMT and Senescence map has the canonical pathway together with details of the RAF1 phosphorylation events (Figure 1d). Additionally, RAF1 from the Regulated Cell Death map is included to represent the potential difficulty of connecting the RAS-RAF-MEK-ERK events to other maps in cases when the state of this protein is not clearly defined (Figure 1b). RAF1 is represented in at least three different ways that would make it difficult to merge and reuse these fragments: RAF1 with no states defined (Figure 1b); RAF1 with one state being shown (Figures 1a and 1c); and RAF1 with six specific phosphorylation sites shown (Figure 1d). Another issue is representing lumped (“generic”) species of MEK and ERK that represent all isoforms (Figures 1a-1c) versus explicitly representing each specific MEK1, MEK2, ERK1 and ERK2 isoforms (Figure 1d).

**Figure 1.**
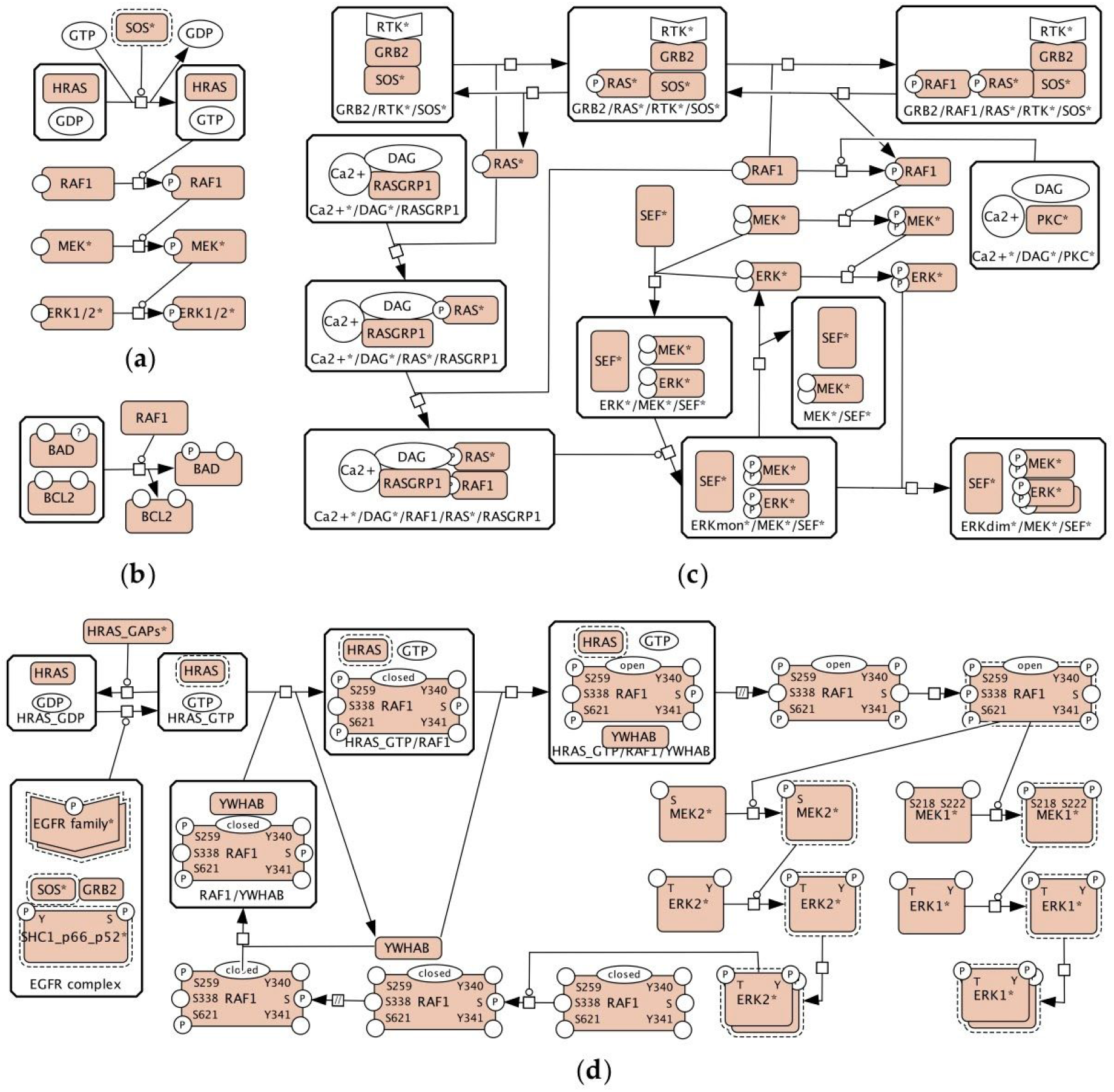
The representation of the RAS-RAF-MEK-ERK pathway in the Atlas of Cancer Signalling Networks [19]: (**a**) The RAS-RAF-MEK-ERK part of the Adaptive Immunity, the Innate Immunity and the Cancer-Associated Fibroblasts maps; (**b**) RAF1 from the Apoptosis module of the Regulated Cell Death map; (**c**) The MAPK part of the Cell Survival map; (**d**) The RAS-RAF-MEF-ERK part of the EMT and Senescence map.

The PANTHER database offers three versions of the RAS-RAF-MEK-ERK pathway in three different maps: two canonical cascades with different levels of detail in the number of phosphorylation sites in Figures 2a and 2b, similarly to Figure 1a, and one version with RAS-RAF-MEK complex formation in Figure 2c that is similar to the map in Figure 1c. “Generic” ERK and MEK are used in all cases, and specific RAF1 is shown in Figure 2b, while “generic” RAF is shown in Figures 2a and 2c.

**Figure 2.**
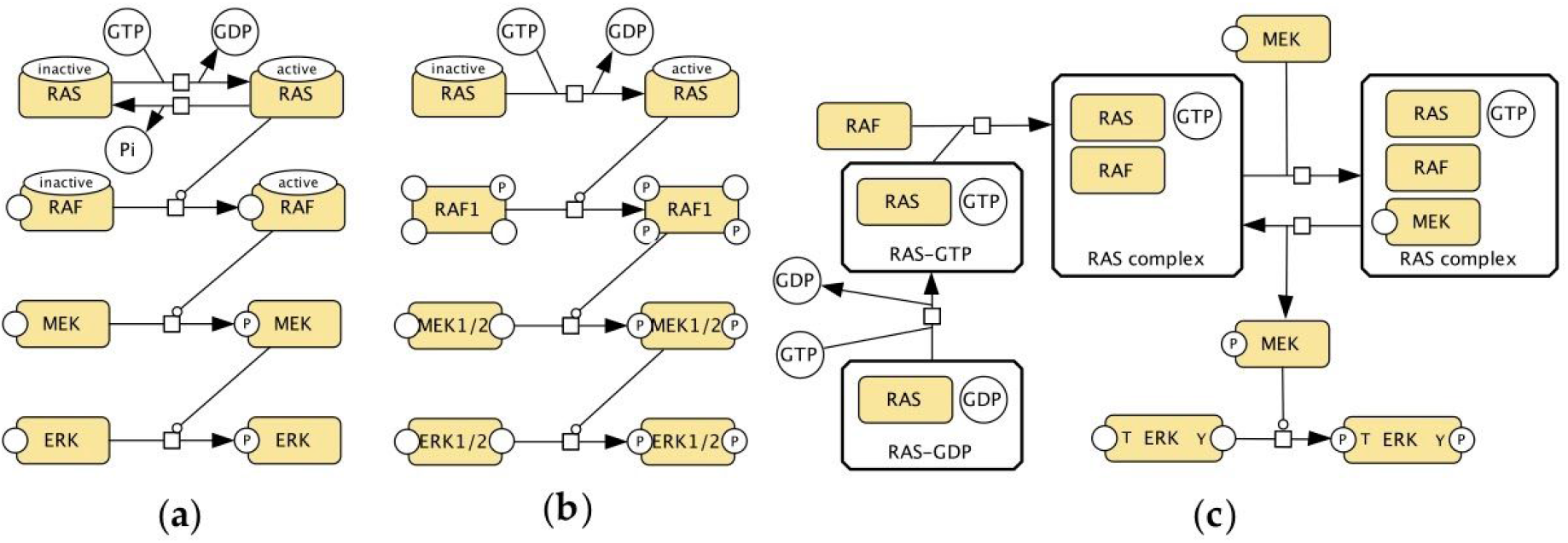
The representation of the RAS-RAF-MEK-ERK pathway in the PANTHER pathway database [20]: (**a**) Part of the Interleukin Signalling Pathway (Pathway:P00036); (**b**) Part of the FGF Signalling Pathway (Pathway:P00021); (**c**) Part of the B Cell Activation Pathway (Pathway:P00010).

Figure 3a shows events inferred from the Reactome RAF/MAP Kinase Cascade map and Figure 3b is a redrawn fragment of the Reactome RAF-independent MAPK1/3 activation map. The canonical representation of the activation of RAF/MEK/ERK monomers (Figures 1a, 2a, 2b) is not found in Reactome. The representation of phosphorylation states of MEK and ERK proteins is consistent in these two maps.

**Figure 3.**
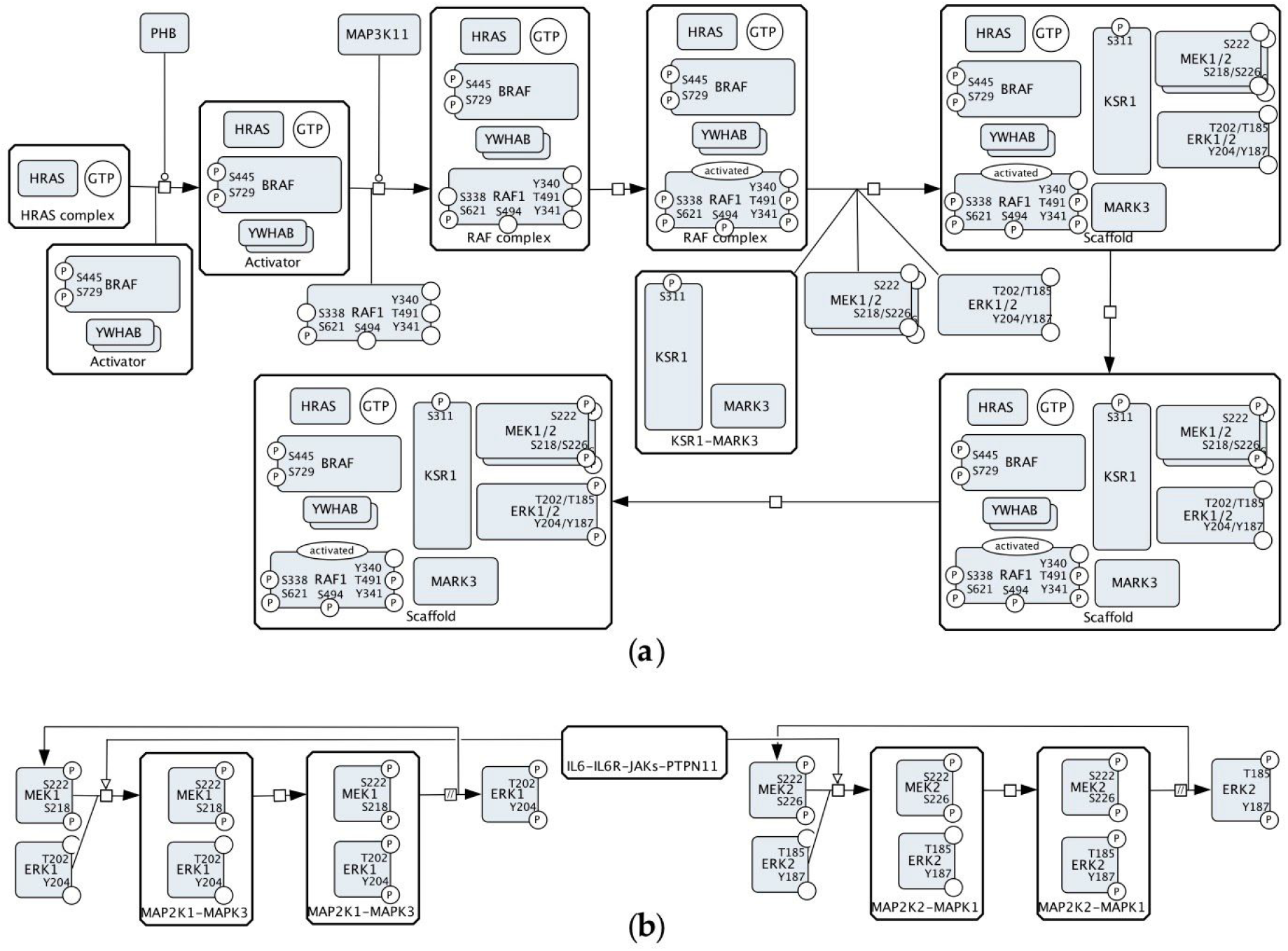
The inferred CellDesigner’s view of MEK-ERK activation events from the Reactome pathway database (https://reactome.org) [21,22]: (**a**) The ERK activation cascade from the RAF/MAP Kinase Cascade map (Pathway: R-HSA-5673001); (**b**) Part from RAF-independent MAPK1/3 Activation map (Pathway:R-HSA-110056).

Reactome has a richer annotation in comparison to other resources: for example, on the RAF/MAP Kinase Cascade map, “generic” RAF includes BRAF, RAF1 and ARAF. On the other hand, it is not always clear how to transform “generic” representation in Reactome into an SBGN-compatible version in CellDesigner. For example, the MEK1/2 entity (Figure 3a) is described in Reactome as “MAPK2K homo/heterodimers” (MEK homo/heterodimers) and has three members: MEK1 homodimer, MEK2 homodimer and MEK1-MEK2 heterodimer. The resulting combinations lead to significantly more extended visualisation (Supplementary Figure S2).

In summary, ACSN, PANTHER, and Reactome contain multiple partially compatible versions of the MAPK pathway: 1) a canonical version of the cascade (Figures 1a, 2a and 2b); 2) a more detailed version with small complexes (Figures 2c and 3b); 3) and a complex regulation via scaffold and adaptor proteins (Figures 1c, 1d and 3a). Phosphorylations sites of RAF1, MEK1, MEK2, ERK1 and ERK2 proteins are represented in different ways with an exception of the Reactome database. “Generic” RAF, MEK and ERK are often used to simplify the representation.

To ensure composability, the components need to be designed in such a way that they are also not conflicting with other parts of the map and it is possible to easily upgrade and replace them with an alternative version. Within the ACSN, the representations of the RAS-RAF-MEK-ERK cascade are parts of larger maps, and as soon as these different representations are harmonised, composability would be ensured and one representation can be replaced by another. The same principle would work for Reactome and PANTHER. The ACSN also sometimes reuses the same component (ERK subnetwork): for example, the representation of Figure 1a is reused in the Adaptive Immunity, the Innate Immunity and the Cancer-Associated Fibroblasts maps. Isolated relevant proteins within other maps, as, for example, “stateless” RAF1 in Figure 1b, need to be identified and aligned with the RAS-RAF-MEK-ERK subnetwork. The same is true about the versions of the PANTHER and Reactome databases. Additionally, Reactome already reuses some of its components and employs submaps as entities of the network. A submap in Reactome is a node that represents a pathway and has a meaning similar to a reusable composable component. For example, the “RAF/MAP kinase cascade” appears as a single-node element on the ERK/MAPK Targets map (Pathway:R-HSA-198753), the FRS-mediated FGFR1 Signalling (Pathway:R-HSA-5654693) and the SHC-mediated Cascade of the FGFR3 Signalling (Pathway:R-HSA-5654704); and double-clicking on this element leads to the RAF/MAP Kinase Cascade map (Pathway:R-HSA-5673001). Despite the reuse of subnetworks within Reactome, there are still different incompatible versions of the RAS-RAF-MEK-ERK pathway and the corresponding compatibility challenges to address. For example, the L1CAM Interactions map (Pathway:R-HSA-373760) contains an entity of MEK1 phosphorylated at S218, S222, T286 and T292 and another entity of MEK1 phosphorylated at T286 and T292 while, normally, activated MEK1 is phosphorylated at S218 and S222.

## 3. Discussion

In terms of reusability, the maps in Figures 1-3 show different versions of events, and a single consensus version is not available. Without additional investigation of the literature, it is not clear which version is the best to reuse, for instance, in a new disease map. There are methodological issues related to the harmonisation of the curation styles and the granularity chosen for capturing events.

This section describes the main barriers to reuse and composition identified during this work and recommendations for dealing with these issues.

### Inconsistent or incompatible descriptions of generic and specific entities and events

By “generic” here we mean an entity that represents a group of proteins. Specific in this context is a particular protein which can be identified in UniProt. If used, the molecular meaning of “generic” entities need to be defined more precisely. For example, the specific proteins lumped into a “generic” entity should be annotated. This enables more compact and modular descriptions of maps. On the other hand, there needs to be a clear molecular description of these groups which is compatible with the description of individual entities and the individual events shown. Examples of conflicting representations: “generic” MEK vs. specific MEK1 and MEK2; and “generic” ERK vs. specific ERK1 and ERK2 (Figure 1a/1c vs. Figure 1d). The use of specific entities is advisable because it allows exact identification of the entities: MEK1 (MAP2K1, UniProt:Q02750), MEK2 (MAP2K2, UniProt:P36507), ERK1 (MAPK3, UniProt:P27361) and ERK2 (MAPK1, UniProt:P28482). It is also important for describing phosphorylations sites when needed. For example, ERK1 is phosphorylated at T202 and Y204 while ERK2 is phosphorylated at T185 and Y187. Automatic identification of such cases is possible via the corresponding queries with the following manual check and semi-automatic replacement of “generic” entities with specific ones.

### Ambiguous or incompatible descriptions of the states of proteins

The rule used for state variables in the provided examples is referred to as “once a variable, always a variable”. Once introduced, a state variable must be applied to all entities of the same protein on a map, even if this state variable is not affected by the represented processes. This rule is enforced in CellDesigner by design, and within the same diagram, once introduced, all state variables are displayed. That means that merging two diagrams is not possible if state variables of the same protein are handled differently. Examples of conflicting representations are shown in Figure 1: RAF1 with no state variables vs. RAF1 with one state variable vs. RAF1 with seven state variables. Such cases need to be harmonised if the module is meant to be reusable and compatible with other parts of the network. Automatic identification and update of all related proteins is feasible but would require manual verification.

### Ambiguous or incompatible descriptions of large complexes

Signalling events include the formation of complexes. The lack of information about such complexes or the lack of standards for describing complexes leads to curators describing them inconsistently. Curators can choose to visualise the corresponding signalling as split events and use smaller complexes, for example, to be able to avoid combinatorial explosion issues when each modification would necessarily lead to the multiplication of similar complexes as shown in Supplementary Figure S2.

### Propagation of alternative variants of the same module

Controversially, while aiming at having a minimal set of reusable and composable modules, we have to consider the necessity of keeping alternative variants of the same component. We need to distinguish between 1) different possible ways to convey the same mechanisms, 2) new levels of complexity introduced, often with more molecules included, and 3) different conditions, cell types or organisms described. Figures 1a, 2a-c and 3b show the same pathway, and they should be merged into one reusable version. Figure 1b and 3a, on the other hand, show different regulatory mechanisms and versioning based on those mechanisms would be beneficial. Mutations can modify a pathway, and that might require a new version (see, for example, an alternative route for mutated protein in the RAF/MAP Kinase Cascade, Pathway:R-HSA-5673001). Another anticipated reason for an alternative view is the possible difference in various cell types under different conditions. A possible solution is re-factoring variants of pathways into a) a “consensus” component and b) deviations from this consensus view. Were possible, maps could be composed of those “consensus” components plus different context-specific deviations would be possible where needed. Propagation of alternative representations assumes developing the corresponding criteria and manual selection of versions to keep within a system.

We believe that solutions to these issues would make ACSN, PANTHER, and Reactome pathways qualitatively more reusable and composable. Methodological improvements should include: a) more explicit curation of the molecular meaning of lumped “generic” species and particular specific proteins; b) standards for describing complexes; c) new ways of describing variants of pathways - consensus pathways with their deviations. This way it is possible to maintain an advanced resource design that would consist of reusable and composable components.

## 4. Methods

To evaluate the reusability and composability of map components, we searched for a pathway with both repeated and different representations among the maps of three different databases: the ACSN, the PANTHER database, and Reactome. We aimed at a signalling pathway since it is the only type of pathways present in the maps of all three databases. To narrow down the list of candidates, we first identified automatically all catalysed protein phosphorylation processes that were repeated among one or more maps of each database, as described below.

For each database, we first stored all maps into a Neo4j graph database (https://neo4j.com). We used stonpy (https://github.com/Adrienrougny/stonpy) to store the maps of the ACSN and PANTHER database and employed the available Neo4j database for Reactome [22] (https://reactome.org/dev/graph-database).

We then queried all catalysed phosphorylation processes using Cypher, the query language for Neo4j. For each database, we used two queries: one for querying phosphorylation processes of free proteins and one for proteins that belong to a complex (Supplementary Table S2). Results of the two queries were obtained in the form of a list of quadruplets: <name of the kinase>, <name of the target>, <phosphorylation site>, <name of the map>. We then grouped the results by the three first values (<name of the kinase>, <name of the target>, <phosphorylation site>) to count the occurrence of each catalysed phosphorylation process, and discarded those that were not repeated. We obtained a list of repeated triplets and associated each triplet with their occurrence in each map, for each database (Supplementary Tables S3-S5).

We then reviewed the lists obtained for the three databases manually in order to find repeated catalyzed phosphorylation processes that would additionally be represented differently among maps. We found that it was the case for the processes of the RAS-RAF-MEK-ERK pathway, and selected this pathway as a suitable candidate to illustrate the issue of composability in the context of cancer research.

Finally, we manually isolated the processes involved in the RAS-RAF-MEK-ERK for six pathways of the ACSN (Figure 1), three pathways of the PANTHER database (Figure 2) and three pathways of Reactome (Figure 3). The selected fragments were then copied (ACSN), redrawn (PANTHER) or reconstructed (Reactome) in CellDesigner, modified for making them visually comparable (for example, association and dissociation glyphs are replaced by a generic process glyph) and additionally manually laid out for better readability.

## 5. Conclusions

Automatic analysis of the ACSN, PANTHER and Reactome databases followed by a manual review of the RAS-RAF-MEK-ERK events demonstrated challenges in reusability and composability. The offered observations and conclusions will be applicable to other pathway resources a well.

Applying such design principles as modularity, reusability and composability is a promising direction for managing the complexity of molecular process networks. Because it is easier to evaluate smaller modules, improve or replace them, reusable and composable components are likely to be more trustworthy and robust. Also, composable components would be more impactful since they could be reused by others. Indirectly, this reuse could lead to more trusted content. If many investigators review and use a component, this gives some confidence that the component is an accurate description of the biology.

We anticipate that if these ideas are adopted, this could lead to a natural improvement of reusable pathway resources. Since a minimal set of versions is discussed, it would allow evolving them in a more focused and controllable way. This could contribute to reducing the number of redundant efforts in pathway biology and to enabling faster development of needed disease models. We are optimistic that development and adoption of the required technologies will enable not only the faster development of maps but more comprehensive and more informative maps that can guide the understanding of the disease and the identification of potential drug targets.

To advance on the topic of reusability and composability and to make it easier for many groups to find information about this signalling cascade, we propose the **RAS-RAF-MEK-ERK Pathway Challenge** (http://sbgn.org/openchallenge) aiming to create a comprehensive up-to-date description of this pathway and offer a minimal set of versions that can be employed in various pathway reconstructions and disease models.

## Supplementary Materials

Table S1: Selected features of the maps from the Atlas of Cancer Signalling Network, PANTHER and Reactome databases, Table S2: Cypher queries used for identifying all catalyzed phosphorylation processes of the pathways of the ACSN, PANTHER and Reactome, Table S3: The analysis of repeated triplets in the ACSN, Table S4: The analysis of repeated triplets in PANTHER, Table S5: The analysis of repeated triplets in Reactome, Figure S1: BRAF signalling component from the ACSN, Figure S2: an alternative version of Figure 3a from Reactome.

## Author contributions

Conceptualisation, AM, AR, JRK, JSR, MO; methodology, AR, AM; investigation, AM, AR; writing - original draft preparation, AM; writing - review and editing, all authors; visualization, AM; project coordination, AM; funding acquisition, RS. All authors have read and agreed to the published version of the manuscript.

