## Supplementary Figure S1 for "Reusability and composability in process description maps: RAS-RAF-MEK-ERK signalling"

**Supplementary Figure S1.** The BRAF fragment of the MAPK module from the Cell Survival map of the Atlas of Cancer Signalling Networks (https://acsn.curie.fr) [Kuperstein 2015 PMID:26192618].

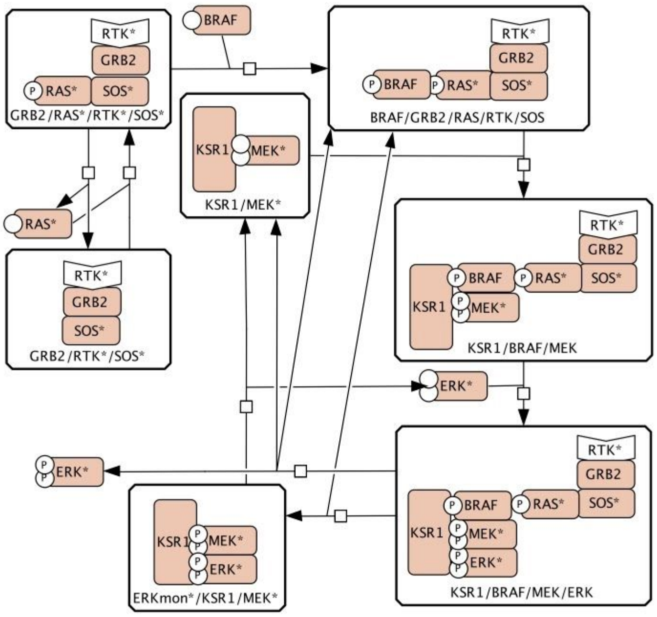
