## Supplementary Figure S2 for "Reusability and composability in process description maps: RAS-RAF-MEK-ERK signalling"

**Supplementary Figure S2**. An inferred CellDesinger's view alternative to Figure 3a - the RAF/MAP Kinase Cascade map (Pathway:R-HSA-5673001) from the Reactome pathway database (https://reactome.org) [Jassal 2020 PMID:31691815; Fabregat 2018 PMID:29377902].

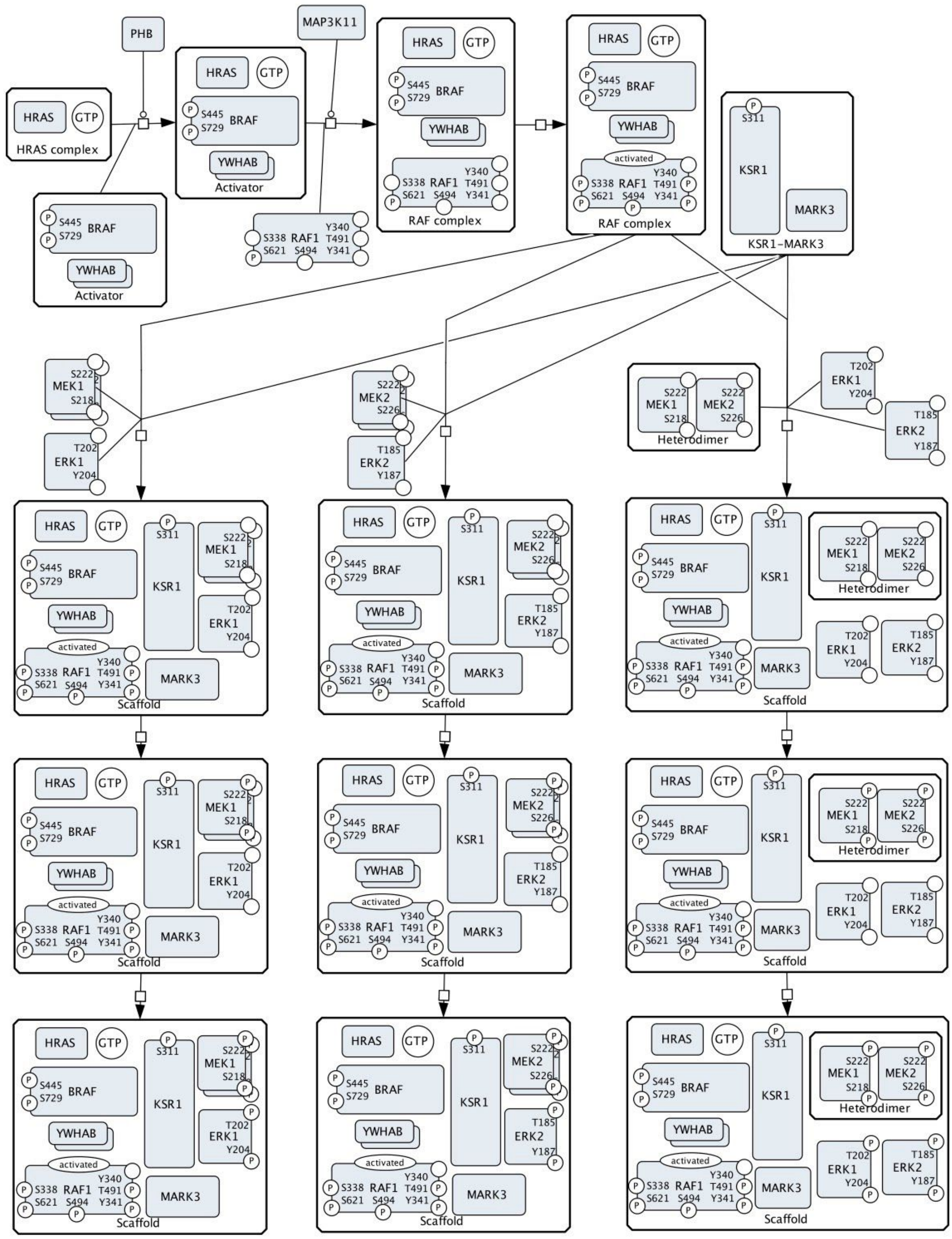
