## Supplementary Table S1 for "Reusability and composability in process description maps: RAS-RAF-MEK-ERK signalling"

**Supplementary Table S1.** Selected features of the maps from the Atlas of Cancer Signalling Network, PANTHER and Reactome databases.

| **Features** | **ACSN** | | | **PANTHER** | | | **Reactome** | | |
|---|---|---|---|---|---|---|---|---|---|
| | **1a** | **1c** | **1d** | **2a** | **2b** | **2c** | **3a** | **3b** | **3c** |
| Identifiers provided for all proteins (UniProt) | ✓ | ✓ | ✓ | ✓ | ✓ | ✓ | ✓ | ✓ | ✓ |
| Identifiers provided for all metabolites (ChEBI) | ✓ | ✓ | ✓ | ✓ | ✓ | ✓ | ✓ | ✓ | ✓ |
| Each protein modification is described properly in SBGN-compatible format | ✓ | ✓ | ✓ | ✓ | ✓ | ✓ | ✓ | ✓ | ✓ |
| Each complex composition is reflected in its content in SBGN-compatible format | ✓ | ✓ | ✓ | ✓ | ✓ | ✓ | ✓ | ✓ | ✓ |
| There are no “mixed” diagrams (no elements of the Reduced Notation of CellDesigner are used) | ✓* | ✓* | ✓* | ✓ | ✓ | ✓ | ✓ | ✓ | ✓ |
| Available formats: CellDesigner | ✓ | ✓ | ✓ | ✓ | ✓ | ✓ | ✗ | ✗ | ✗ |
| Available formats: SBGN-ML 0.2 | ✓ | ✓ | ✓ | ✗ | ✗ | ✗ | ✓ | ✓ | ✓ |
| Available formats: SBGN-ML 0.3** | ✗ | ✗ | ✗ | ✗ | ✗ | ✗ | ✗ | ✗ | ✗ |
| Available formats: SBML*** | ✓ | ✓ | ✓ | ✓ | ✓ | ✓ | ✓ | ✓ | ✓ |
| Available formats: BioPAX Level 2 | ✗ | ✗ | ✗ | ✗ | ✗ | ✗ | ✓ | ✓ | ✓ |
| Available formats: BioPAX Level 3 | ✗ | ✗ | ✗ | ✓ | ✓ | ✓ | ✓ | ✓ | ✓ |

*Rare cases
**Recently published version with the support for annotation and colours
***With CellDesigner’s XML available it is possible to export the content in SBML Level 1 Version 2 - SBML Level 2 Version 4
